# Retrospective Evaluation of the Eye Irritation Potential of Agrochemical Formulations

**DOI:** 10.1101/2023.06.23.543016

**Authors:** Neepa Choksi, Andreia Latorre, Shadia Catalano, Arthur Grivel, James Baldassari, Janaina Pires, Marco Corvaro, Mariana Silva, Maryanne Ogasawara, Monique Inforzato, Priscila Habe, Rosana Murata, Stefan Stinchcombe, Susanne Kolle, W. Masinja, Gisele Perjessy, Amber Daniel, David Allen

## Abstract

While multiple *in vitro* eye irritation methods have been developed and adopted as OECD health effects test guidelines, only one is considered a complete replacement for the *in vivo* rabbit eye test for classification and labelling for eye irritation hazard. Additionally, for most of the adopted methods there is a data gap on the applicability to predict the ocular irritation potential of agrochemical formulations. To overcome this data gap, a retrospective evaluation of 192 agrochemical formulations with *in vivo* (OECD TG 405) or *in vitro* (OECD TG 437, 439, or 492) data was conducted to determine if the *in vitro* methods could accurately assign United Nations Globally Harmonized System for Classification and Labelling of Chemicals (GHS) eye irritation hazard classifications. In addition, for each of the final formulations and their individual components, we also conducted an evaluation of the GHS concentration threshold (CT) approach. The results herein suggest that the four individual methods and GHS CT approach were highly predictive of formulations that would not require classification for eye irritation hazard. Given most agrochemical formulations fall into this category, methods that accurately identify not classified mixtures could significantly reduce the use of animals for this endpoint.

## 1 Introduction^1^

The *in vivo* rabbit eye test (commonly known as the Draize eye test) has been used for more than 75 years to assess the irritation and corrosion potential of substances and formulations that may come into contact with the eye (Draize et al., 1944) (Table S1). This test relies upon subjective assessments of reversibility and damage that is produced as a result of test substance exposure. While historically used, the assay is subject to interobserver and animal variations which are likely responsible for the irreproducibility of this test method, particularly for mild and moderate eye irritants (Luechtefeld et al. 2016). Therefore, for both scientific and ethical reasons, alternatives to the *in vivo* rabbit eye test have been considered to replace or reduce animal use for this endpoint. For example, the United Nations Globally Harmonized System for Classification and Labelling of Chemicals (GHS) includes a concentration threshold (CT) approach for classifying eye irritation. This approach uses the skin and eye irritation classification on the components to assign an eye hazard classification if the sum of the components exceeds specific concentration limits. Applying such an approach could reduce animal use by waiving tests conducted on the end-use product.

Several *in vitro* eye irritation test methods have been validated for identification of seriously eye damaging substances, and for identifying chemicals as “not classified” (NC) or not requiring signal words based on decision criteria for the GHS hazard classification and labelling system. Some of these methods have been adopted as test guidelines issued by the Organisation for Economic Co-operation and Development (OECD), which assist in acceptance of data across countries and reduce repeated testing (OECD, 2020, 2018, 2019). Only in 2022 the first method considered complete replacements that are capable of identifying the full spectrum of eye irritation i.e. to predict GHS Categories 1, 2 and “not classified” chemicals has been adopted as OECD Test Guideline (TG) 492B (OECD 2022). However, the applicability of this test method has not yet been evaluated for agrochemical formulations. Furthermore, none of the adopted test methods is able to subcategorize Category 2A (moderate irritants) and Category 2B (mild irritants).

Agrochemical formulations are typically mixtures (multi-constituent substances, under GHS definition) composed of one or more active ingredients combined with one or more co-formulants to optimize activity and enhance delivery of the active ingredient(s) (U.S. Environmental Protection Agency, 2022). In August 2019, based on new guidance approximately 2,000 formulations registered in Brazil were reclassified according to the GHS hazard classification system (Brazil 3a Diretoria Gerencia-Geral de Toxicologia, 2019). Almost 70% of the formulated products registered in Brazil were classified as NC for eye irritation according to GHS classification system. Identification of non-animal methods that could be used to classify NC formulations could lead to a significant reduction in the number of animals needed for evaluation of eye irritation potential.

Due to the complex nature of mixtures and formulations, these substances are not typically included as reference chemicals in test method validation efforts. *In vitro* eye irritation testing of agrochemical formulations has reported discordant results with classifications based on *in vivo* rabbit studies. These studies have partially been limited based on the number of formulations evaluated (Kolle et al., 2017a; Settivari et al., 2016). An exception is the Reconstructed Human Cornea-Like Epithelium (RhCE) method EpiOcular™ Eye Irritation Test, which has been shown to be predictive to identify agrochemical formulations non-irritant to the eye (Kolle et al., 2015). Hence, agrochemical formulations have been inclusively mentioned in the respective OECD TG 492.

Herein we evaluate the possibility to reduce the use of animals for eye irritation testing of agrochemical formulations, by involving retrospective evaluations of existing but previously unpublished data. We describe a retrospective evaluation of the usefulness and limitations of multiple *in vitro* ocular irritation test methods and the GHS CT approach to assign hazard classification and labelling for eye irritation potential for agrochemical formulations.

## 2 Materials and Methods

### 2.1 Agrochemical Formulation Data Submissions

Data for agrochemical formulations were submitted by CropLife Brasil companies (BASF, Bayer Crop Science, Corteva Agriscience, FMC, Ihara, Ourofino, Sumitomo Chemical and Syngenta) to Inotiv, a contract research organization that created a database inclusive of all submitted data and conducted the analyses presented herein. Availability of (1) associated historical rabbit data or GHS ocular irritancy classification from test method conducted according to OECD TG 405, and (2) results from an *in vitro* eye irritation method or (3) results from the GHS CT approach were required for inclusion in the analyses (OECD, 2021; United Nations, 2021). No additional animal testing was conducted for this evaluation. Formulation types were obtained from individual study reports or provided directly by the submitting companies and were compliant with the CropLife International guidance (CLI, 2017).

### 2.2 Test Methods

*In vitro* results were available from at least one of three test methods: Bovine Corneal Opacity and Permeability (BCOP); Isolated Chicken Eye (ICE); and the RhCE. All methods were conducted according to their respective OECD TGs (i.e., OECD TG 437 for BCOP, OECD TG 438 for ICE, and OECD TG 492 for RhCE) (OECD, 2020, 2018, 2019), under Good Laboratory Practices (GLP) or GLP-like conditions. General information regarding conduct of each test method protocol was provided by the submitting companies.

### 2.3 Data Analyses

Submitted *in vitro* assay and *in vivo* rabbit test data were compiled into a single spreadsheet. Classifications based on *in vitro* test method results, using the decision criteria included in their respective OECD TG, were compared to GHS classifications derived from historical *in vivo* rabbit test data to evaluate concordance. A result was judged to be concordant when classification based on *in vitro* results agreed with classification based on *in vivo* rabbit data. A result was judged to be discordant when classification based on *in vitro* results did not agree with classification based on *in vivo* rabbit data. For formulations that were identified as “no prediction can be made” by the *in vitro* method (see Table S2), the results were identified as discordant with the *in vivo* prediction since a definitive classification could not be assigned based on the *in vitro* result.

For each *in vitro* test method, classifications were compared to *in vivo* classifications using a bottom-up and top-down approach (Scott et al., 2010). Specifically, for the top-down approach, the ability of an *in vitro* test method to identify serious eye damage (i.e., *in vitro* and *in vivo* methods independently predicted a formulation should be classified as Category 1) was evaluated (Category 1 *vs* Categories 2 and Not Classified). Likewise, for the bottom-up approach, the ability of an *in vitro* test method to identify formulations that do not require eye irritation labels (i.e., *in vitro* and *in vivo* methods independently predicted a formulation should be classified as NC) was evaluated (Categories 1 and 2 *vs* Not Classified).

Data were also evaluated based on formulation type to determine if there were any specific types associated with particularly high or low concordance. These analyses were first done based on the formulation types provided by the submitting companies, and then based on three more broad formulation groups: solid, liquid water-based, and liquid solvent-based. A summary of the formulation types included in this evaluation and the grouping scheme used for this analysis is shown in Table S3.

### 2.4 GHS Concentration Threshold Approach

To determine if the eye irritation potential of a formulation could be predicted based on the collective toxicity of its individual components, a standardized worksheet was developed. Each company performed the CT approach as described in the GHS for classification of mixtures using the worksheet (see Figure 1). Each company obtained the eye and skin irritation classifications of the individual components and assessed the pH for each of their respective formulations. None of the formulations had extreme pH (13 < pH < 2.5) or skin corrosion (generally rare for agrochemical formulations) which was deemed useful in prediction of the *in vivo* rabbit outcomes (Kolle et al., 2017b; Corvaro et al., 2017). In most cases, these data were obtained from external sources (i.e., material safety data sheets from the chemical supplier; from Table 3 of Annex VI to the CLP Regulation; REACH registration data from ECHA) (UK Health and Safety Executive, 2021), according to the workflow (see Figure 1). After obtaining the classification of each component, the relevance of these components was determined (see Figure 1). When relevant, the component was included in the calculation, which was performed following the GHS CT approach criteria (Figure S1). The hazard classification based on the threshold approach was compared to that obtained from the *in vivo* rabbit eye test for the entire formulation and concordance determined.

**Figure 1.**
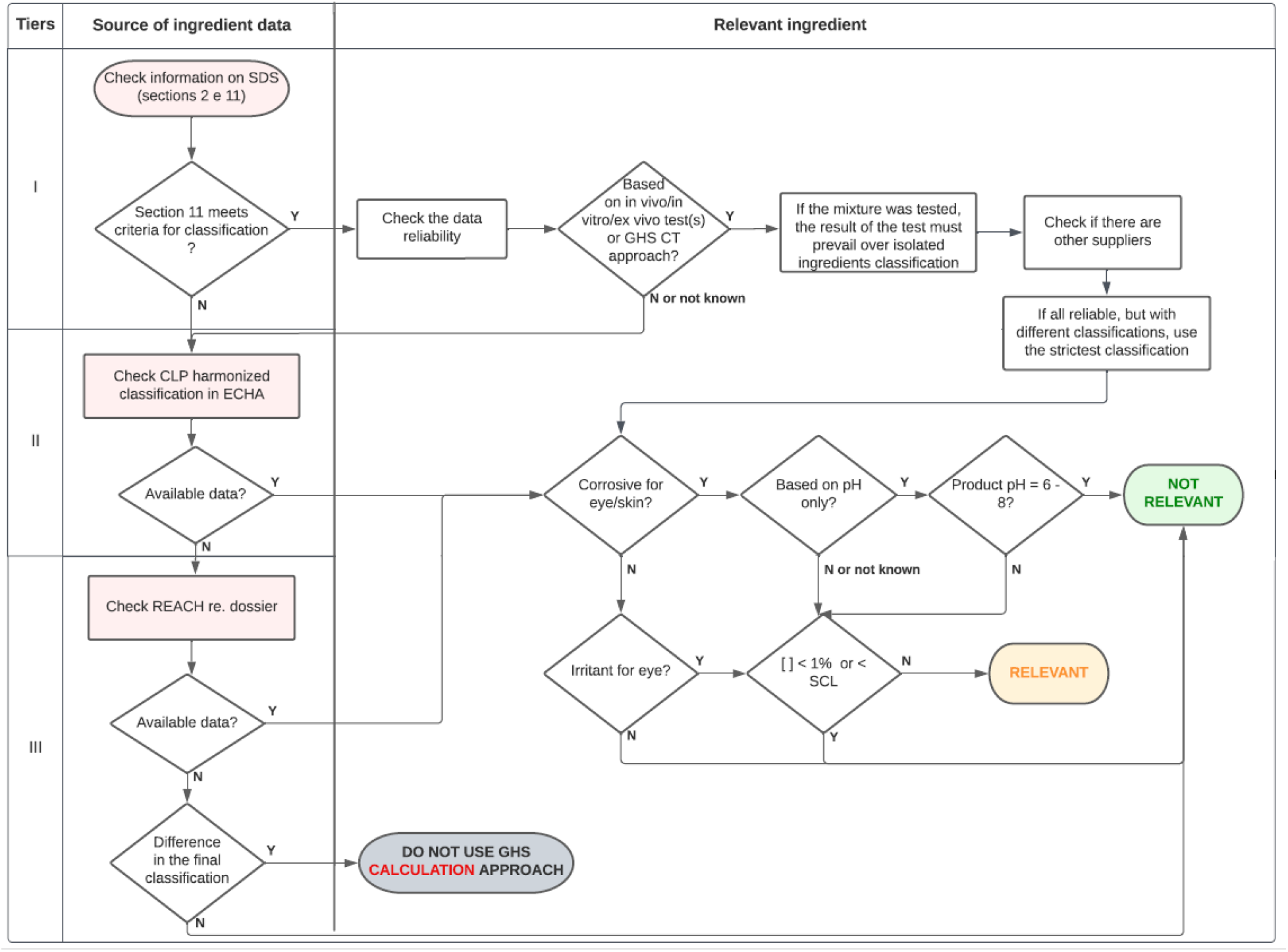
Workflow to verify reliable information on eye irritation of each compound within a formulation, which was used to perform the concentration threshold approach as described in the GHS for classification of mixtures.

## 3. Results

### 3.1 Submitted Data from CropLife Brazil (CLB) Companies

Data on 192 formulations with retrospective *in vivo* data were received from eight different agrochemical companies to evaluate the GHS CT approach; of those 158 had corresponding *in vitro* data (Table 1). No overlap in tested formulations was noted between companies, based on the information provided. Fifteen formulations, submitted by a single company, were tested in two *in vitro* test methods; 14 were tested using the BCOP and RhCE test methods and one formulation was tested using RhCE and ICE test method. The remaining formulations were tested in a single test method. Review of the supporting *in vivo* data showed that all the formulations that were classified as Category 1 were assigned based on the presence of a persistent effect on day 21 in at least one tested animal (data described in each test result). Among the 192 submitted formulations with retrospective *in vivo* data, 11% were Category 1, 18% were Category 2, and 71% were not classified.

**Table 1.** Submitted Test Data

| Company | BCOP | RhCE | ICE | GHS CT approach | Total Substances |
| --- | --- | --- | --- | --- | --- |
| BASF | 21* | 71*<sup>#</sup> | 1<sup>#</sup> | 77 | 77 |
| Bayer | - | - | 29 | 28 | 29 |
| Corteva | - | 10 | - | 10 | 10 |
| FMC | - | - | - | 20** | 20 |
| Ihara | 4 | - | - | 0 | 4 |
| Ourofino | 7 | - | - | 7 | 7 |
| Sumitomo | - | - | - | 15** | 15 |
| Syngenta | 5 | - | 25 | 30 | 30 |
| **Total** | 37 | 81 | 55 | 187 | 192 |

\*14 formulations were tested in the BCOP and the RhCE assays
<sup>#</sup>1 formulation tested in the RhCE and ICE assays
\*\*Only *in vivo* data submitted for the GHS CT approach evaluation.

### 3.2 Concordance Analysis

#### 3.2.1 Bovine Corneal Opacity and Permeability (BCOP) assay

The overall concordance rate for BCOP *vs*. GHS *in vivo* following the top-down and bottom-up approaches were 70% (26/37) and 49% (18/37), respectively (Tables S4 and 2). In the top-down approach, all formulations identified as Category 1 eye irritants based on *in vivo* response, were classified as “No stand-alone prediction can be made” by the BCOP test method. Further review showed that for 78% (7/9) of the formulations classified as Category 1, the classification was based on the response in a single animal. The remaining two formulations showed responses in more than a single animal. For all the Category 1 formulations, the observed response was present at the end of the evaluation period (i.e., 21 days).

In both analysis methods, the negative predictivity was > 70%. This indicates that a negative response by the BCOP method is likely to be identified as a negative by the *in vivo* test method.

#### 3.2.2 Isolated Chicken Eye (ICE)

The overall concordance rates for ICE vs. GHS *in vivo* following the top-down and bottom-up approaches were 85% (47/55) and 78% (43/55), respectively (Tables S5 and 2). In the top-down approach, all formulations identified as Category 1 eye irritants based on *in vivo* response, were classified as “No stand-alone prediction can be made” by the ICE test method. Further review showed that all eight formulations classified as Category 1 were based on a response in a single animal; 75% (6/8) of these formulations were tested in only one animal. Additionally, all Category 1 classifications were based on the persistence of a response until the final day of the evaluation period. In both analysis methods, the negative predictivity was ≥ 85%.

#### 3.2.3 Reconstructed Human Cornea-Like Epithelium (RhCE)

Compared to the BCOP and ICE *in vitro* methods, the decision criteria for RhCE only enable the evaluation of a bottom-up analysis approach. The overall concordance of the RhCE method was 77% (62/81) (Tables S6 and 2). Further review showed that for 33% (2/6) of the formulations classified as Category 1, the classification was based on the response in a single animal. The remaining four formulations showed responses in more than a single animal. All formulations identified as Category 1 or Category 2 using results from the *in vivo* test were identified as “No stand-alone prediction can be made” by the RhCE method. The negative predictivity of the method was 100% (46/46).

#### 3.2.4 Performance of the GHS Concentration Threshold Approach

The GHS category for 187 agrochemical formulations was (1) provided based on *in vivo* test results with the final formulation, and (2) calculated based on the GHS CT approach. Within class concordance was calculated for each GHS category, with the notable exception that GHS category was not subcategorized into Category 2A or 2B because those categories are not provided by the calculation results. Within category concordance was highest for Categories 1 (62% [13/21]) and Not Classified (58% [77/133]) whereas Category 2 concordance was only 27% [9/33] (Table S7).

Among the Category 1 formulations, the majority of the discordant results (87.5% [7/8]) were formulations predicted by the GHS CT approach as Category 2. Among the Category 2 formulations (n=33) most of the discordant formulations (19/24) were overpredicted by the GHS CT approach as Category 1. The remaining five discordant formulations were underpredicted by the GHS CT approach as NC. Finally, among the discordant not classified formulations (56/133), 21 were formulations overpredicted by the GHS CT approach as Category 1, and 35 were overpredicted by the GHS CT approach as Category 2. However, of the 83 formulations predicted as not classified by the GHS CT approach, almost all (92% [77/83]) are concordant with the *in vivo* classification; 5/6 discordant formulations are GHS Category 2B *in vivo*. Therefore, a GHS calculation that results in no classification appears to be a high confidence prediction.

However, much like the *in vitro* methods, applying a bottom-up and top-down approach greatly improved test method performance, particularly with regard to negative predictivity which was 93–94%.

### 3.3 Evaluation Based on Formulation Type

Results from evaluation of concordant or discordant prediction rates using the BCOP test method based on specific formulation types were limited by the number of formulations of a particular type. As shown in Table S8, the discordance rate for the WG/WDG formulation type was 100% (6/6). The discordance rates for EC and SC formulation types were 70% (7/10) and 72% (8/11), respectively. A closer look at these observed discordances did not show a consistent pattern of over- or under-prediction when compared to the *in vivo* rabbit classification. For example, as shown in Table S9, all five of the EC formulations that were classified as Category 1 based on *in vivo* responses were identified as “No stand-alone prediction can be made” by the BCOP method. Comparatively, for *in vivo* Category 2 formulations, a single formulation was under-classified when compared to the *in vitro* classification. Finally, for *in vivo* Category NC formulations, 2/3 EC formulations were discordant. Similar lack of pattern based on formulation type was observed with the ICE and RhCE results (data not shown).

To interrogate the concordance predictions further, we grouped the formulations according to three formulation-type groups: solids, liquid water-based, and liquid solvent-based. As shown in Table 3, *in vitro* and *in vivo* concordance rates varied greatly between formulation-type groups and classification analysis. For both the BCOP and ICE test method, the top-down analysis approach yielded the highest overall *in vitro* concordance rates for solids and liquid water-based formulations. Comparatively, higher concordance rates were observed using the bottom-up approach for liquid solvent-based formulations. Negative predictivity rates for all formulation-type groups and analysis methods were greater than 50%, with most rates ≥ 75%.

## 4 Discussion and Conclusions

Agrochemical formulations are typically mixtures composed of one or more active ingredients combined with one or more “inert” constituents to optimize activity and enhance delivery of the active ingredient(s) (U.S. Environmental Protection Agency, 2022). Prior to registration and use, companies submit to regulatory authorities’ information about the acute, subchronic, and chronic effects potentially caused by their active ingredients. Usually, information on acute toxicity properties including skin and eye irritant properties (colloquially named as the “6-pack”) is required for the end-use products (agrochemical formulations). Although to date several non-animal methods have been adopted to assess ocular irritancy as OECD TGs, to date only the RhCE method EpiOcular™ Eye Irritation Test (EIT) has been described to predict agrochemical formulations sufficiently well for non-irritants (Kolle et al., 2015). Animal studies are generally conducted to fulfill regulatory requirements. As noted above, almost 70% of the formulations registered in Brazil until 2019 did not require classification for eye irritation according to GHS. Identification of methods that could be used to classify NC formulations could lead to a significant reduction in the number of animals tested for evaluation of eye irritation potential.

Luechtefeld and colleagues showed that the likelihood of replicate *in vivo* rabbit eye tests yielding the same classification is < 50% for substances which are initially identified as mild or moderate irritants (Luechtefeld et al., 2016). Reproducibility concerns, in conjunction with animal welfare concerns and regulations banning or restricting animal testing, have led to the development and evaluation of non-animal test methods (Oliveira et al., 2015).

In the present study, we conducted a retrospective evaluation using agrochemical formulations that had existing *in vivo* and *in vitro* eye irritation data (and were not included in previously published data sets).

The three *in vitro* methods included in this evaluation were BCOP, RhCE, and ICE; all three methods are included in OECD Health Effects TGs (OECD, 2020, 2018, 2019). These methods represent a variety of domains of applicability and coverage of key biological mechanisms of irritation. For example, the BCOP and ICE methods provide a full-thickness model to assess corneal effects. The RhCE model allows for measurement of cytotoxicity, a critical event in the irritation pathway, in cells from the species (i.e., human) of interest (Clippinger et al., 2021). None of the methods (all with a single short-term exposure), however, have been described to detect persistent effects, which is known to be the main trigger of classification as UN GHS Category 1 for eye irritation of agrochemical formulations (Adriaens et al., 2014; Barroso et al., 2017). In fact, in the present dataset all the 21 formulations assigned Category 1 (11% [21/192]) were based on the presence of a persistent effect on day 21 in at least one tested animal.

Although binary concordance between *in vitro* and GHS threshold approach and *in vivo* results using a top-down or bottom-up approach was moderate for all four test methods evaluated, the negative predictivity rate (i.e., that the probability that a negative result obtained by the *in vitro* test method or GHS threshold approach was also identified as a negative by the *in vivo* method), for the four methods was > 85% (Table 2B). The high negative predictivity rate indicates that a formulation identified as not requiring hazard classification based on an *in vitro* test method or GHS threshold approach, is not likely to be an eye irritant. Coupled with the recognized variability of the *in vivo* test method, one might consider the *in vitro* methods or GHS threshold approach to be sufficiently predictive of such effects for regulatory applications (Weil and Scala, 1971; Earl et al., 1997; York and Steiling, 1998; Luechtefeld et al., 2016).

**Table 2.** Concordance Analysis

| Test Method | n | Concordance |  | False Positive Rate |  | False Negative Rate |  | Positive Predictivity |  | Negative Predictivity |  |
| --- | --- | --- | --- | --- | --- | --- | --- | --- | --- | --- | --- |
|  |  | % | No. | % | No. | % | No. | % | No. | % | No. |
| BCOP (Top-down) | 37 | 70 | 26/37 | 7 | 2/28 | 100 | 9/9 | 0 | 0/2 | 74 | 26/35 |
| ICE (Top-down) | 55 | 85 | 47/55 | 0 | 0/47 | 100 | 8/8 | - | - | 85 | 47/55 |
| GHS CT approach (Top-down) | 187 | 74 | 139/187 | 24 | 40/166 | 38 | 8/21 | 25 | 13/53 | 94 | 126/134 |

**B. Bottom-up analyses**
| Test Method | n | Concordance |  | False Positive Rate |  | False Negative Rate |  | Positive Predictivity |  | Negative Predictivity |  |
| --- | --- | --- | --- | --- | --- | --- | --- | --- | --- | --- | --- |
|  |  | % | No. | % | No. | % | No. | % | No. | % | No. |
| BCOP (Bottom-up) | 37 | 49 | 18/37 | 75 | 18/24 | 8 | 1/13 | 40 | 12/30 | 86 | 6/7 |
| ICE (Bottom-up) | 55 | 78 | 43/55 | 22 | 8/36 | 21 | 4/19 | 65 | 15/23 | 88 | 28/32 |
| RhCE (Bottom-up) <sup>a</sup> | 81 | 77 | 62/81 | 29 | 19/65 | 0 | 0/16 | 46 | 16/35 | 100 | 46/46 |
| GHS CT approach (Bottom-up) | 187 | 67 | 125/187 | 42 | 56/133 | 11 | 6/54 | 54 | 56/104 | 93 | 77/83 |

**Table 3.** Concordance Analysis Based on Formulation Group

| Test Method | Formulation Group | n | Concordance |  | False Positive Rate |  | False Negative Rate |  | Positive Predictivity |  | Negative Predictivity |  |
| --- | --- | --- | --- | --- | --- | --- | --- | --- | --- | --- | --- | --- |
|  |  |  | % | No. | % | No. | % | No. | % | No. | % | No. |
| BCOP (Top-down) | Solids | 8 | 75 | 6/8 | 0 | 0/6 | 100 | 2/2 | - | - | 75 | 6/8 |
| BCOP (Top-down) | Water-Based | 17 | 82 | 14/17 | 7 | 1/15 | 100 | 2/2 | 0 | 0/1 | 88 | 14/16 |
| BCOP (Top-down) | Solvent-Based | 12 | 50 | 6/12 | 14 | 1/7 | 100 | 5/5 | 0 | 0/1 | 55 | 6/11 |
| ICE (Top-down) | Solids | 11 | 91 | 10/11 | 0 | 0/10 | 100 | 1/1 | - | - | 91 | 10/11 |
| ICE (Top-down) | Water-Based | 31 | 97 | 30/31 | 0 | 0/30 | 100 | 1/1 | - | - | 97 | 30/31 |
| ICE (Top-down) | Solvent-Based | 13 | 54 | 7/13 | 0 | 0/6 | 100 | 6/6 | - | - | 54 | 7/13 |
| GHS CT approach (Top-down) | Solids | 31 | 64 | 20/31 | 34 | 10/29 | 50 | 1/2 | 9 | 1/10 | 95 | 19/20 |
| GHS CT approach (Top-down) | Water-Based | 89 | 85 | 76/89 | 14 | 12/87 | 50 | 1/2 | 8 | 1/13 | 99 | 75/76 |
| GHS CT approach (Top-down) | Solvent-Based | 50 | 60 | 30/50 | 42 | 15/36 | 36 | 5/14 | 38 | 9/24 | 81 | 21/26 |

**B. Bottom-up analyses**
| Test Method | Formulation Group | n | Concordance |  | False Positive Rate |  | False Negative Rate |  | Positive Predictivity |  | Negative Predictivity |  |
| --- | --- | --- | --- | --- | --- | --- | --- | --- | --- | --- | --- | --- |
|  |  |  | % | No. | % | No. | % | No. | % | No. | % | No. |
| BCOP (Bottom-up) | Solids | 8 | 25 | 2/8 | 100 | 6/6 | 0 | 0/2 | 25 | 2/8 | - | - |
| BCOP (Bottom-up) | Water-Based | 17 | 47 | 8/17 | 64 | 9/14 | 0 | 0/3 | 25 | 3/12 | 100 | 5/5 |
| BCOP (Bottom-up) | Solvent-Based | 12 | 67 | 8/12 | 75 | 3/4 | 13 | 1/8 | 70 | 7/10 | 50 | 1/2 |
| ICE<br>(Bottom-up) | Solids | 11 | 64 | 7/11 | 40 | 2/5 | 33 | 2/6 | 67 | 4/6 | 60 | 3/5 |
| ICE<br>(Bottom-up) | Water-Based | 31 | 81 | 25/31 | 18 | 5/28 | 33 | 1/3 | 29 | 2/7 | 96 | 23/24 |
| ICE<br>(Bottom-up) | Solvent-Based | 13 | 85 | 11/13 | 33 | 1/3 | 10 | 1/10 | 90 | 9/10 | 67 | 2/3 |
| RhCE<br>(Bottom-up)<sup>a</sup> | Solids | 15 | 47 | 7/15 | 57 | 8/14 | 0 | 0/1 | 11 | 1/9 | 100 | 6/6 |
| RhCE<br>(Bottom-up)<sup>a</sup> | Water-Based | 42 | 88 | 37/42 | 13 | 5/38 | 0 | 0/4 | 44 | 4/9 | 100 | 33/33 |
| RhCE<br>(Bottom-up)<sup>a</sup> | Solvent-Based | 24 | 75 | 18/24 | 46 | 6/13 | 0 | 0/11 | 65 | 11/17 | 100 | 7/7 |
| GHS CT<br>approach<br>(Bottom-up) | Solids | 31 | 58 | 18/31 | 54 | 12/22 | 11 | 1/9 | 40 | 8/20 | 91 | 10/11 |
| GHS CT<br>approach<br>(Bottom-up) | Water-Based | 89 | 73 | 65/89 | 27 | 21/78 | 27 | 3/11 | 28 | 8/29 | 95 | 57/60 |
| GHS CT<br>approach<br>(Bottom-up) | Solvent-Based | 50 | 70 | 35/50 | 70 | 14/20 | 3 | 1/30 | 67 | 29/43 | 86 | 6/7 |

<sup>a</sup>RhCE method does not have decision criteria to allow for a top-down evaluation (i.e., no decision criteria for GHS Category 1). Therefore, only a bottom-up analysis was conducted.

The formulation type may have an impact on whether a specific formulation produces an *in vitro* result that is concordant with the *in vivo* response. To assess the impact of formulation type, individual and group formulations were evaluated for concordance. The limited number of individual formulation types tested by each *in vitro* test method limited the conclusions that could be made by this evaluation. However, the higher prevalence of end-use products not requiring classification for eye irritant is different when considering groups of formulation types, being the higher prevalence among liquid water-based formulations. So, this information can be considered in the weight of evidence analysis to evaluate the prediction of alternative testing strategies.

To increase the available data to conduct these analyses, specific formulation types were combined into one of three formulation groups: solid, liquid solvent-based, or liquid water-based. The top-down analysis approach yielded the highest overall *in vitro* concordance rates for solids and liquid water-based formulations for BCOP and ICE test methods. Comparatively, higher concordance rates were observed using the bottom-up approach for solvent-based formulations. Overall, the negative predictivity rates for all formulation-type groups and analysis methods were > 50%, with most rates ≥ 75%. As noted above, the high negative predictivity rate indicates that a formulation identified as not requiring hazard classification based on an *in vitro* test method, is not likely to be an eye irritant.

A recent publication characterized the available *in vivo* and *in vitro* test methods with respect to their relevance to human ocular anatomy, anticipated exposure scenarios, and the mechanisms of eye irritation/corrosion (Clippinger et al., 2021). The *in vitro* methods were shown to be at least as relevant to the effects observed in the human eye when compared to the rabbit eye.

Overall, these studies suggest that *in vitro* test methods, as well as the GHS CT approach, could be used to classify agrochemical formulations as not needing classification. All formulation group types appeared to be equivalently evaluated by the evaluated test methods. Given that approximately 70% of registered agrochemical formulations were assigned the hazard classification of not classifiable for eye irritation according to GHS classification system, the use of *in vitro* methods would lead to a significant reduction in the number of animals tested for evaluation of eye irritation potential.

Accordingly, Figure 2 provides a proposed decision workflow that could be used for the assessment of eye irritation/corrosion potential of agrochemical formulations using new approach methodologies (NAM).

**Figure 2.**
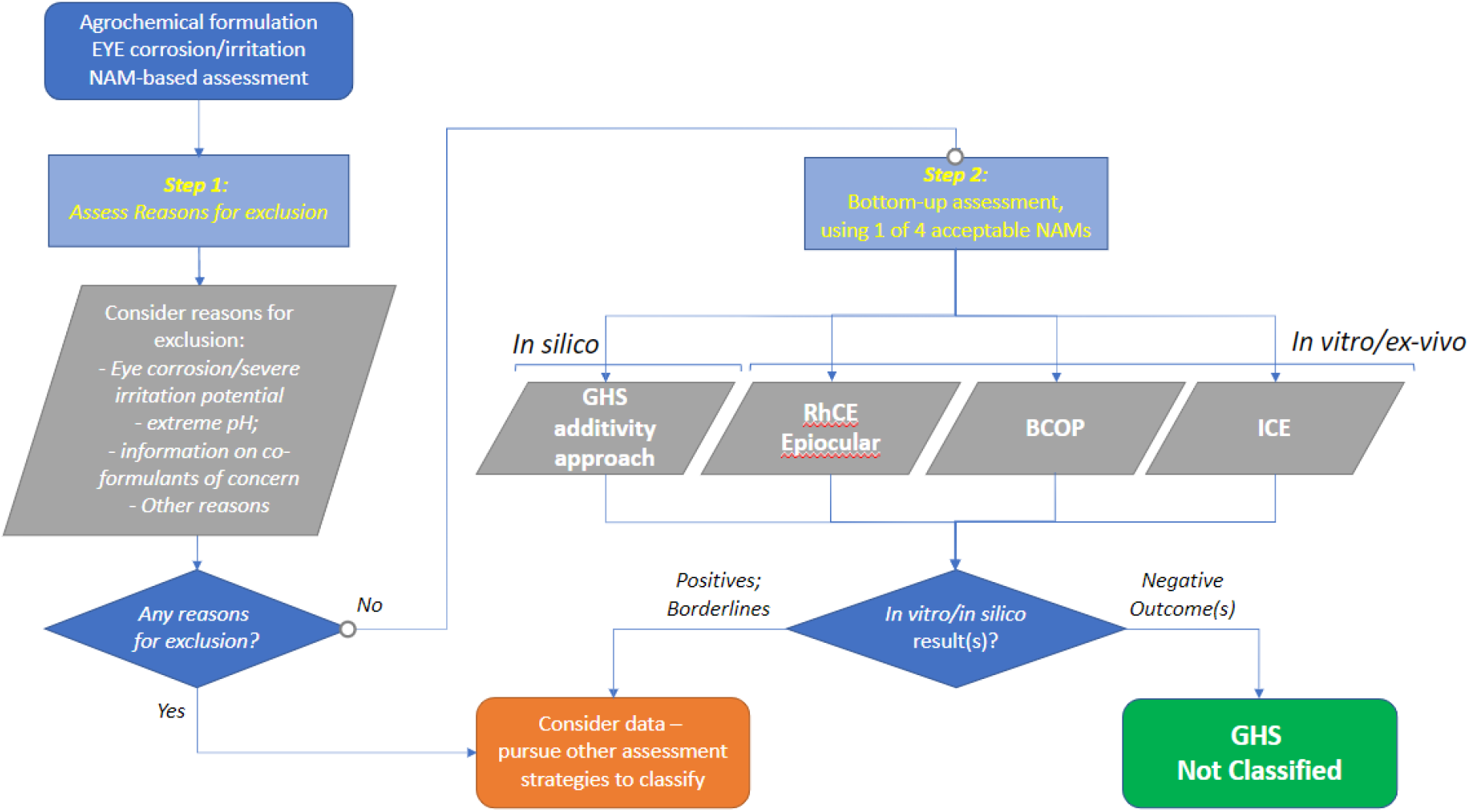
Proposed decision flowchart for assessment of agrochemical formulation eye irritation/corrosion potential, using NAM-based assessment. After initial considerations on high potential for corrosivity (Step 1), the suggested approach is to start with a bottom-up assessment (Step 2), using one of the four methods for which negative predictivity was deemed acceptable based on our retrospective review. In this step, one or more approaches can be used depending on the applicability (i.e., availability of information on all co-formulants for the GHS calculation; compatibility with the test system for the three *in vitro* assays).

While the analyses presented herein do not provide a solution for the long-awaited complete replacement for animal use in eye irritation testing, they do provide promise for a path toward reducing animal use through possible test waivers (i.e., using the GHS threshold approach) and rationale for filling data gaps to better identify a potential Defined Approach for eye irritation testing without using animals at all.

## Abbreviations

GHS: United Nations Globally Harmonized System for Classification and Labelling of Chemicals
CT: concentration threshold
NC: not classified
RhCE: Reconstructed Human Cornea-Like Epithelium
BCOP: Bovine Corneal Opacity and Permeability
ICE: Isolated Chicken Eye
TG: Test Guideline
NAM: new approach methodologies

## CRediT authorship contribution statement

The authors confirm contribution to the paper as follows: **study conception and design:** Andreia Latorre and Shadia Catalano; **data collection:** Andreia Latorre, Arthur Grivel, James Baldassari, Janaina Pires, Marco Corvaro, Mariana Silva, Maryanne Ogasawara, Monique Inforzato, Priscila Habe, Rosana Murata, Shadia Catalano, Stefan Stinchcombe, Susanne Kolle and William Masinja; **software:** Gisele Perjessy**; analysis and interpretation of results:** Neepa Choksi and Dave Allen; **draft manuscript preparation:** Neepa Choksi; **results review:** Andreia Latorre**; writing review:** Andreia Latorre, Arthur Grivel, Gisele Perjessy, James Baldassari, Janaina Pires, Marco Corvaro, Mariana Silva, Maryanne Ogasawara, Monique Inforzato, Priscila Habe, Rosana Murata, Shadia Catalano, Stefan Stinchcombe, Susanne Kolle, and William Masinja; **editing**: Dave Allen; **supervision:** Gisele Perjessy; **project administration:** Gisele Perjessy and Dave Allen. All authors reviewed and approved the final version of the manuscript.

## Data availability statement

Data are available upon request from the corresponding author

## Acknowledgements

The authors thank Elizabeth Farley-Dawson, PhD, for her editorial review of this manuscript.

