## Supplemental material for "Retrospective Evaluation of the Eye Irritation Potential of Agrochemical Formulations"

**Article Type:** Research Article

[1]Inotiv, Research Triangle Park, NC, USA; [2]BASF SA, São Paulo, SP, Brazil; [3]Corteva Agriscience, Newark, DE, USA; [4]Bayer SAS, Sophia Antipolis, France; [5]FMC Corporation, Philadelphia, PA, USA; [6]Ouro Fino Química S.A, Ribeirão Preto, SP, Brazil; [7]Corteva Agriscience Italia S.p.A, Rome, Italy; [8]Iharabras S.A. Indústrias Químicas, Sorocaba, SP, Brazil; [9]FMC Corporation, Campinas, SP, Brazil; [10]Syngenta Crop Protection, Greensboro, USA; [11]Sumitomo Chemical Brasil Indústria Química S.A., São Paulo, SP, Brazil; [12]Bayer SA, São Paulo, SP, Brazil;[13]BASF SE, Ludwigshafen, Germany;[14]Syngenta Ltd., Bracknell, United Kingdom; [15]CropLife Brasil, São Paulo, SP, Brazil.

*Current affiliation for William Masinja is Intertek, Farnborough, UK; for Mariana Silva is Corteva Agroscience, Alphaville, SP, Brazil; for Neepa Choksi is ToxStrategies, RTP, NC;

### SUPPLEMENTAL INFORMATION

**Table S1. GHS Ocular Irritation Classification Systems**

| GHS Category | Classification Criteria[a] |
|---|---|
| 1 | Effects on the cornea, iris, or conjunctiva that are not expected to reverse or do not fully reverse within 21 days |
| 2 | Effects on the cornea, iris, or conjunctiva that fully reverse within 21 days |
| NC | No effects are produced, or minimal effects observed that do not lead to classification |

Abbreviations: NC = not classified; PPE = personal protective equipment

[a] Category 1 GHS classification is applied when a substance produces either (a) mean corneal opacity score ≥ 3 or iritis score ≥ 1.5 (over Days 1, 2, and 3) in at least two animals or (b) a score > 0 on Day 21. A Category 2 classification is applied when a substance produces either (a) mean corneal opacity or iritis score ≥ 1 or (b) conjunctival redness score ≥ 1 (over Days 1, 2, and 3) in at least 2 animals.

**Table S2. *In Vitro* Test Method Decision Criteria**

| Test Method | Calculations/Endpoints Evaluated | Category 1 | No Prediction Can Be Made | Not Classified |
|---|---|---|---|---|
| BCOP | IVIS = mean opacity + (15 × mean permeability)* | IVIS > 55 | 3 < IVIS ≤ 55 | ≤ 3 |
| RhCE | Mean tissue viability | No criteria available | ≤ 60% | > 60% |
| ICE | Combination of corneal swelling, corneal opacity, and fluorescein retention endpoint scores | 3 x IV<br>2 x IV, 1 x III<br>2 x IV, 1 x II<br>2 x IV, 1 x I<br>CO = 3 at 30 min (≥2 eyes)<br>CO = 4 at any time point (≥2 eyes)<br>Severe loosening of the epithelium (≥1 eye) | Combinations not noted in "Not Classified" or "Category 1" hazard classifications | 3 x I<br>2 x I, 1 x II<br>2 x II, 1 x I |

Abbreviations: CO = corneal opacity, IVIS = *in vitro* irritancy score

*Calculation referent the Opacitometer OP-KIT* and Duratec

| Sum of ingredients classified as: | Concentration triggering classification of a mixture as: | |
|---|---|---|
| | **Serious eye damage** | **Eye irritation** |
| | **Category 1** | **Category 2/2A** |
| Skin Category 1 + Eye Category 1[a] | ≥ 3% | ≥ 1% but < 3% |
| Eye Category 2 | | ≥ 10%[b] |
| 10 x (skin Category 1 + eye Category 1)[a] + eye Category 2 | | ≥ 10% |

[a]If an ingredient is classified as both skin Category 1 and eye Category 1 its concentration is considered only once in the calculation;

[b]A mixture may be classified as eye Category 2B when all relevant ingredients are classified as eye Category 2B.

**Figure S1.** Globally Harmonized System of Classification and Labelling of Chemicals (GHS) concentrations of ingredients of a mixture classified as skin Category 1 or Category 1 or 2 that would trigger classification of the mixture as hazardous to the eye (Category 1 or 2) (United Nations, 2021)**.**

**Table S3. Formulation-Type Groups (adapted from CLI, 2017)**

| Formulation group | Individual formulation types |
|---|---|
| Solid | GR (granule)<br>RB (bait, ready for use)<br>WG/WDG (water dispersible granule)<br>WP (wettable powder) |
| Liquid, water-based | AE (aerosol dispenser)[a]<br>CS (capsule suspension)<br>FS (flowable concentrate for seed treatment)<br>SC (suspension concentrate)<br>SD (suspension concentrate for direct application)<br>ZC (mixed formulation of CS and SC formulation types)<br>SL (soluble concentrate)[a] |
| Liquid, solvent-based | AE (aerosol dispenser)[b]<br>DC (dispersible concentrate)<br>EC (emulsifiable concentrate)<br>EW (emulsion, oil in water)<br>GL (emulsifiable gel)<br>OD (oil dispersion)<br>SE (suspo-emulsion)<br>SL (soluble concentrate)[b] |

(a) total organic solvent and surfactant concentration < 30% in the formulation; the threshold of 30% was empirically derived based on the distribution of the formulations tested.

(b) total organic solvent and surfactant concentration ≥ 30% in the formulation.

**Table S4.** Confusion Matrices for Top-Down (A) and Bottom-Up (B), and Complete Cross-Reference for BCOP (C)

**A.**

| | *In Vitro* | |
|---|---|---|
| *In Vivo* | Cat. 1 | Other |
| Cat. 1 | 0 | 9 |
| Other | 2 | 26 |

**B.**

| | *In Vitro* | |
|---|---|---|
| *In Vivo* | Other | NC |
| Other | 12 | 1 |
| NC | 18 | 6 |

**C.**

| | *In Vitro* | | | |
|---|---|---|---|---|
| *In Vivo* | Cat. 1 | NP | NC | Total |
| Cat. 1 | 0 | 9 | 0 | 9 |
| Cat. 2 | 1 | 2 | 1 | 4 |
| NC | 1 | 17 | 6 | 24 |
| Total | 2 | 28 | 7 | **37** |

NC: Not classified; NP: No prediction can be made.

**Table S5.** Confusion Matrices for Top-Down (A) and Bottom-Up (B), and Complete Cross-Reference for ICE (C)

**A.**

| | *In Vitro* | |
|---|---|---|
| *In Vivo* | Cat.1 | Other |
| Cat. 1 | 0 | 8 |
| Other | 0 | 47 |

**B.**

| | *In Vitro* | |
|---|---|---|
| *In Vivo* | Other | NC |
| Other | 15 | 4 |
| NC | 8 | 28 |

**C.**

| | *In Vitro* | | | |
|---|---|---|---|---|
| *In Vivo* | Cat. 1 | NP | NC | Total |
| Cat. 1 | 0 | 7 | 1 | 8 |
| Cat. 2 | 0 | 8 | 3 | 11 |
| NC | 0 | 8 | 28 | 36 |
| Total | 0 | 23 | 32 | **55** |

NC: Not classified; NP: No prediction can be made.

**Table S6.** Confusion Matrices for Bottom-Up (A), and Complete Cross-Reference for RhCE (B)

**A.**

| | *In Vitro* | |
|---|---|---|
| *In Vivo* | NP | NC |
| Other | 16 | 0 |
| NC | 19 | 46 |

**B.**

| | *In Vitro* | | | |
|---|---|---|---|---|
| *In Vivo* | | NP | NC | Total |
| Cat. 1 | | 6 | 0 | 6 |
| Cat. 2 | | 10 | 0 | 10 |
| NC | | 19 | 46 | 65 |
| Total | | 35 | 46 | **81** |

NC: Not classified; NP: No prediction can be made.

**Table S7.** Confusion Matrices for Top-Down (A) and Bottom-Up (B), and Complete Cross-Reference for GHS CT approach (C)

**A.**

| | GHS CT approach | |
|---|---|---|
| *In Vivo* | Cat. 1 | Other |
| Cat. 1 | 13 | 8 |
| Other | 40 | 126 |

**B.**

| | GHS CT approach | |
|---|---|---|
| *In Vivo* | Other | NC |
| Other | 48 | 6 |
| NC | 56 | 77 |

**C.**

| | GHS CT approach | | | |
|---|---|---|---|---|
| *In Vivo* | Cat. 1 | Cat. 2 | NC | Total |
| Cat. 1 | 13 | 7 | 1 | 21 |
| Cat. 2 | 19 | 9 | 5 | 33 |
| NC | 21 | 35 | 77 | 133 |
| Total | 53 | 51 | 83 | **187** |

NC: Not classified; NP: No prediction can be made.

**Table S8.** Concordance Analysis for Individual Formulation Types Tested in the BCOP Test Method

| Formulation Type | Concordant Prediction* | Discordant Prediction |
|---|---|---|
| EC (n=10) | 3 | 7 |
| EW (n=1) | - | 1 |
| FS (n=3) | 1 | 2 |
| OD (n=1) | - | 1 |
| SC (n=11) | 3 | 8 |
| SE (n=2) | 1 | 1 |
| WG/WDG (n=6) | - | 6 |
| WP (n=2) | - | 2 |
| Unknown (n=1) | - | 1 |

Abbreviations: EC = emulsifiable concentrate, EW = emulsion, oil in water, FS = flowable concentrate for seed treatment, OD = oil dispersion, SC = suspension concentrate, SE = suspo-emulsion, WG/WDG = water disposable granule, WP = wettable powder.

* Predictions considered concordant:

*In vivo* Cat. 1 predicted as Cat. 1

*In vivo* Cat. 2A/2B classified as no stand-alone prediction can be made

*In vivo* NC predicted as NC

**Table S9.** Cross-Reference of BCOP Results Based on *In vivo* Category and Formulation Type

| *In Vivo* GHS Category | Formulation Type | BCOP: 1 | BCOP: NP | BCOP: NC |
|---|---|---|---|---|
| 1 | EC | - | 5 | - |
| | EW | - | 1 | - |
| | SC | - | 1 | - |
| | WG/WDG | - | 2 | - |
| 2 | EC | - | 2 | 1 |
| | SC | 1 | - | - |
| NC | EC | 1 | 1 | 1 |
| | FS | - | 2 | 1 |
| | OD | - | 1 | - |
| | SC | - | 6 | 3 |
| | SE | - | 1 | 1 |
| | WG/WDG | - | 4 | - |
| | WP | - | 2 | - |

Abbreviations: EC = emulsifiable concentrate, EW = emulsion, oil in water, FS = flowable concentrate for seed treatment, NC = not classified, NP = no stand-alone prediction can be made, OD = oil dispersion, SC = suspension concentrate, SE = suspo-emulsion, WG/WDG = water disposable granule, WP = wettable powder.
